# Serial dependence is absent at the time of perception but increases in visual working memory

**DOI:** 10.1101/187666

**Authors:** Daniel P. Bliss, Jerome J. Sun, Mark D’Esposito

**Affiliations:** UC Berkeley, Helen Wills Neuroscience Institute, Berkeley, CA, USA; UC Berkeley, Department of Psychology, Berkeley, CA, USA

## Abstract

Recent experiments have shown that visual cognition blends current visual input with that from the recent past to guide ongoing decision making. This serial dependence is tuned to the similarity between consecutive stimuli and appears to exploit the temporal autocorrelation normally present in visual scenes to promote perceptual stability. While these benefits have been assumed, evidence that serial dependence directly alters stimulus perception has been limited. In the present study, we parametrically vary the delay between stimulus and response in a spatial delayed response task to explore the trajectory of serial dependence from the moment of perception into post-perceptual visual working memory. We find that behavioral responses made immediately after viewing a stimulus show evidence of adaptation, but not attractive serial dependence. Only as the memory period lengthens is a blending of past and present information apparent in behavior, reaching its maximum with a memory delay of six seconds. These results dovetail with other recent findings to bolster the interpretation that serial dependence is a phenomenon of mnemonic rather than perceptual processes. We also demonstrate that when leading mathematical models of visual working memory are adjusted to account for this trial-history effect, their fit to behavioral data is substantially improved.

## Introduction

Even in contexts where visual input varies randomly from trial to trial, human observers tend to blend stimuli from previous trials into their representation of the current one, leading to a bias in behavioral reports^1–14^. This smoothing of representations – termed “serial dependence” – is a function of how close successive stimuli are in space^1,15^ and time^1–7,12,14,15^. It is also sensitive to their featural similarity^1–3, 7–10, 12, 14, 16, 17^. Serial dependence has been observed in judgments of orientation^1,8,9^ and location^16,17^, as well as more complex attributes like the identity^2^ and attractiveness^3,5,11^ of human faces. That the bias is observed for such disparate features suggests it may be a universal principle of visual processing, and recent work has sought to demonstrate its adaptiveness^8^: In natural environments – where the input to our eyes is generally very similar from moment to moment^18^ – temporal smoothing would be expected to stabilize perception in the face of noise and occlusion^1,8^.

While the benefits of perceptual stability seem obvious, it is important to note that serial dependence impedes another goal of perception, which is to be sensitive to change. A classic example of how visual perception prioritizes change detection is the tilt after-effect^19^. This illusion (which is a type of adaptation^20^) is the quantitative opposite of serial dependence: Perception of the current moment is repelled away from, rather than merged with, recently processed stimuli – exaggerating differences. Like serial dependence, adaptation spans different types of stimulus features^20–24^. However, unlike adaptation, the attractive bias depends on attention: the observer must attend to each stimulus for serial dependence to occur^1^. Attention is thought to rely on the same neural and psychological mechanisms as working memory^25–31^. Hence, it is possible that whereas adaptation is a phenomenon of visual perception^20–24^ serial dependence arises instead from post-perceptual visual working memory^9,32^. If this were true, stability would operate in parallel with (rather than compete against) change detection, as these functions would be relegated to distinct cognitive systems^9^.

Preliminary efforts have been made to resolve whether serial dependence is perceptual or mnemonic in nature, with mixed results^1,9^. Using a comparison task that minimizes memory demands, one group identified positive serial dependence in a small number of individuals^1^ – in favor of the perceptual account. However, an attempt to replicate this effect with a larger sample size only revealed repulsive adaptation^9^. That is, no attractive serial dependence was observed when memory demands were removed from the same comparison task in the follow-up study, in support of the idea that serial dependence requires working memory^9^. A complementary strategy for clarifying this issue has been to boost memory demands – by increasing the delay between stimulus and response in delayed-estimation tasks – to determine whether this potentiates the attractive bias^9,32^. Traditionally, errors that scale with delay length are interpreted as mnemonic in origin, whereas those that are constant over time are assumed to be tied to the perceptual or motor demands that are also fixed^33^. Over a limited range, the magnitude of the serial dependence effect increases the longer that working memory is active^9,16,17^. Despite this potential connection to working memory, serial dependence has yet to be incorporated into the many mathematical models that have been developed in recent years to fit the dispersion of errors in human memory-guided behavior^34–43^.

In the present study, we investigate temporal smoothing in visual cognition over a wider range of memory delays than has been used in the past. We use a spatial delayed response task, which has been shown to produce serial dependence in non-human primates^16,17^. Previous experiments using delayed response tasks to measure serial dependence have included a visual mask after the stimulus presentation period^1,2,8,9^, as well as a delay period of at least several hundred milliseconds before a response is permitted^1,2,8,9,16,17^, which encourages encoding into working memory and cannot cleanly measure more fragile perceptual representations^44,45^. In our shortest delay condition, we allow participants to respond immediately after stimulus offset, with no mask. From this 0-s baseline, we parametrically increase the delay length up to 10 s. In a separate experiment, we parametrically manipulate the length of the inter-trial interval (ITI). This permits us to assess the decay rate of the trial-history effect in the absence of intervening trials – clarifying its potential functional and biological implementation. Finally, we pursue a novel formal unification of the serial dependence phenomenon with mathematical models of working memory^34,35,37–41^. This sets the stage for future experiments to dissect the neural mechanisms of serial dependence in the context of ongoing research into the organization of the working memory system^32^.

## Results

### Experiment 1: Manipulation of visual working memory delay

Participants completed a spatial delayed response task, depicted in Figure 1. For Experiment 1, the length of the working memory delay period in this task was varied randomly from trial to trial (0, 1, 3, 6, or 10 s). Collapsing across these five delay conditions, we identified serial dependence in the group dataset significantly greater than zero *(p <* 10^−4^, group permutation test; peak-to-peak = 1.67°; bootstrapped 95% confidence interval = [1.48°, 1.85°]). To do this analysis, we measured the magnitude of serial dependence as the peak-to-peak of the curve fit to the pattern of errors across all possible differences between current and previous stimulus location (see Methods). The peak-to-peak is a measure of the maximal pull of responses away from the correct stimulus feature value as a result of this trial-history bias. Previous studies have used similar measures of amplitude to quantify serial dependence^1,2,8,9,16^. No bias was present in the data in the direction of the stimulus on the upcoming trial *(n.s.,* group permutation test; peak-to-peak = −0.14°; bootstrapped 95% confidence interval = [−0.59°, 0.15°]), which supports the conclusion that the dependence of behavior on the previous trial is not due to spurious correlations in the particular randomized sequences of stimuli generated for the subjects^10,14^.

**Figure 1.**
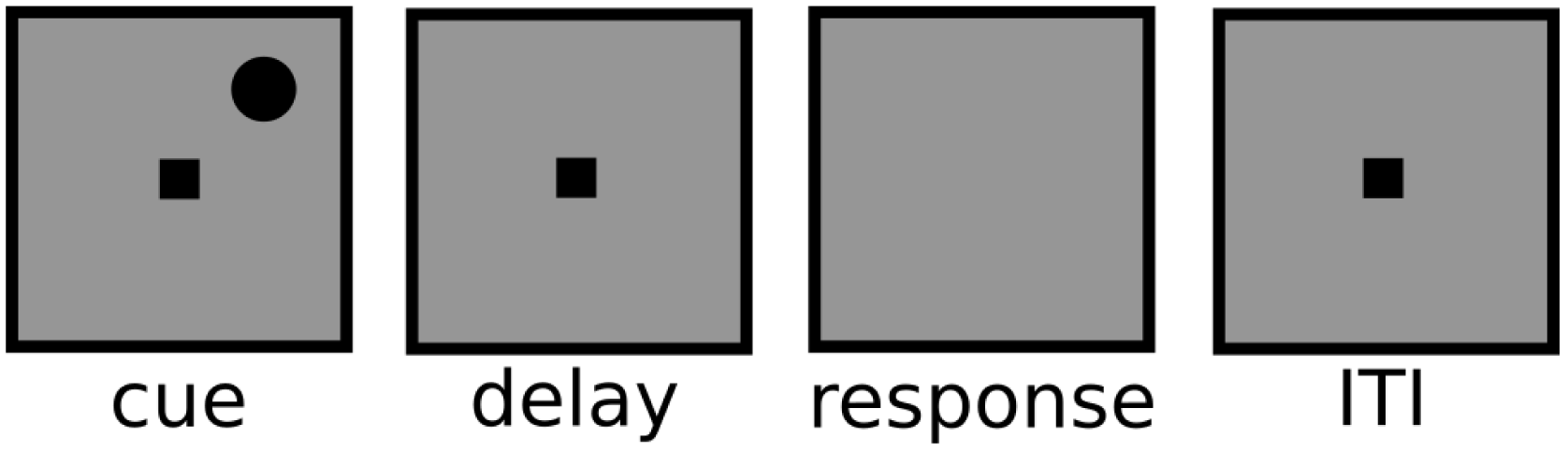
The events in each trial of the generic spatial judgment task used for Experiments 1 and 2 (not to scale, see Methods for exact dimensions). Stimuli were presented in black against a gray background. Participants maintained fixation at the central square whenever it was on the screen (all task stages aside from the response period). Each trial started with the presentation of the cue whose location needed to be remembered for either a variable (Experiment 1) or fixed (Experiment 2) delay. Upon the disappearance of the central square at the end of the delay, the mouse cursor appeared at the exact center of the screen (not shown), and subjects used the mouse to make their response. Responses were not timed. Immediately after the response was made, the fixation square returned for a fixed (Experiment 1) or variable (Experiment 2) inter-trial interval.

Next we examined each of the delays individually. The magnitude of serial dependence across memory delays from 0-10 s is plotted in Figure 2A. When participants reported the location of the stimulus immediately after viewing it, presumably relying at least in part on residual neural activity associated with perception, their responses showed signs of sensory adaptation, an effect that is opposite in direction from serial dependence *(p <* 10^−4^, group permutation test; peak-to-peak = −1.72°; bootstrapped 95% confidence interval = [−2.30°, −1.09°]; Fig. 2B). In contrast, for every other delay tested, serial dependence was significantly greater than zero (all *p <* 0.01, group permutation tests). Moreover, the magnitude of serial dependence increased from 0-1 s *(p <* 10^−4^, group permutation test; peak-to-peak at 1 s = 0.85°; bootstrapped 95% confidence interval = [0.48°, 1.20°]) and again from 3-6 s *(p <* 10^−3^; peak-to-peak at 6 s = 3.37°; bootstrapped 95% confidence interval = [2.88°, 3.84°]) before asymptoting between 6 and 10 s *(n.s.;* peak-to-peak at 10 s = 2.86°; bootstrapped 95% confidence interval = [2.28°, 3.41°]). Serial dependence was numerically strongest in the 6-s condition, shown in Figure 2C. Here, the peak-to-peak is visible as the distance along the y-axis between the maximal and minimal values of the model fit to the data. We note that the asymptote in serial dependence between 6 and 10 s does not correspond to an asymptote in the accumulation of noise in working memory. Consistent with a recent theoretical study and reanalysis of empirical data^?^, we observed a sublinear increase in the variance of responses, which in the case of our data continued up to the 10 s delay (bootstrapped 95% confidence interval at 6 s = [41.28°, 44.92° ], at 10 s = [50.22°, 54.40° ]; Fig. 3).

**Figure 2.**
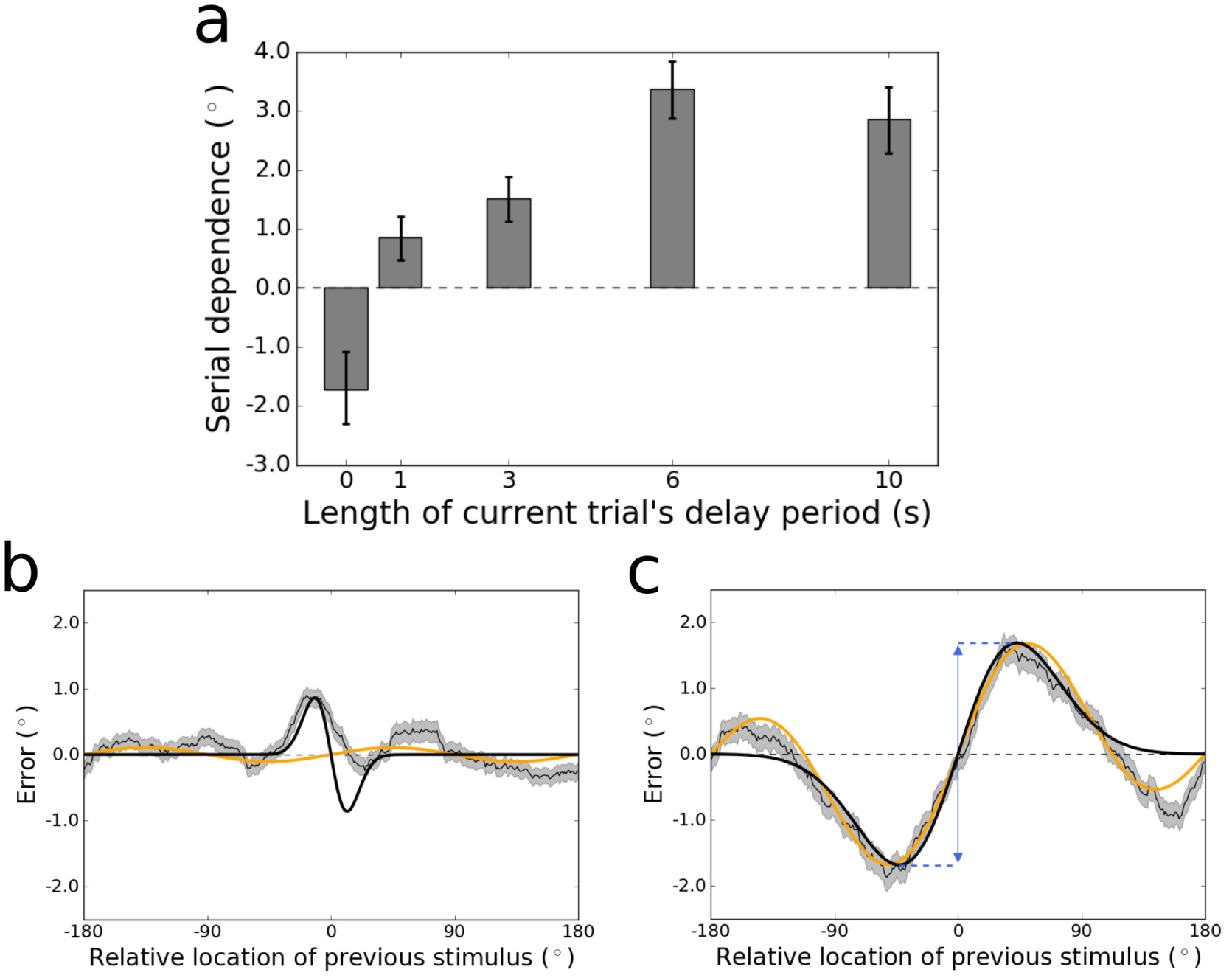
(A) Magnitude of serial dependence in the group data for each delay period tested in Experiment 1. Serial dependence is measured as the peak-to-peak of a least squares fit of the derivative of Gaussian (DoG) tuning function to the data. Error bars represent bootstrapped 95% confidence intervals. The magnitude of the serial dependence increases during the first 6 s of the delay period, and goes from significantly negative (evident of sensory adaptation) to significantly positive between 0 and 1 s. (B) Tuning of serial dependence across all possible angular differences between the current and previous stimulus, for the 0-s delay condition. The thin black line represents the group moving average of response errors, with the standard error in gray shading. The thick black line is the best-fitting DoG curve, and the orange line depicts the best fit of an alternative model – the Clifford model (see Methods) – which cannot capture the pattern of sensory adaptation in this condition. Although its positive and negative peaks are asymmetrical, adaptation is significantly stronger than chance at 0 s, with a peak-to-peak of −1.72^◦^. (C) Tuning of serial dependence for the 6-s delay condition. Here, serial dependence is significantly more positive than chance, and the peak-to-peak of the DoG fit – indicated by the blue double-headed arrow in the figure – is 3.37^◦^. Note that in this condition both the DoG and Clifford models capture the amplitude of the effect equivalently well.

**Figure 3.**
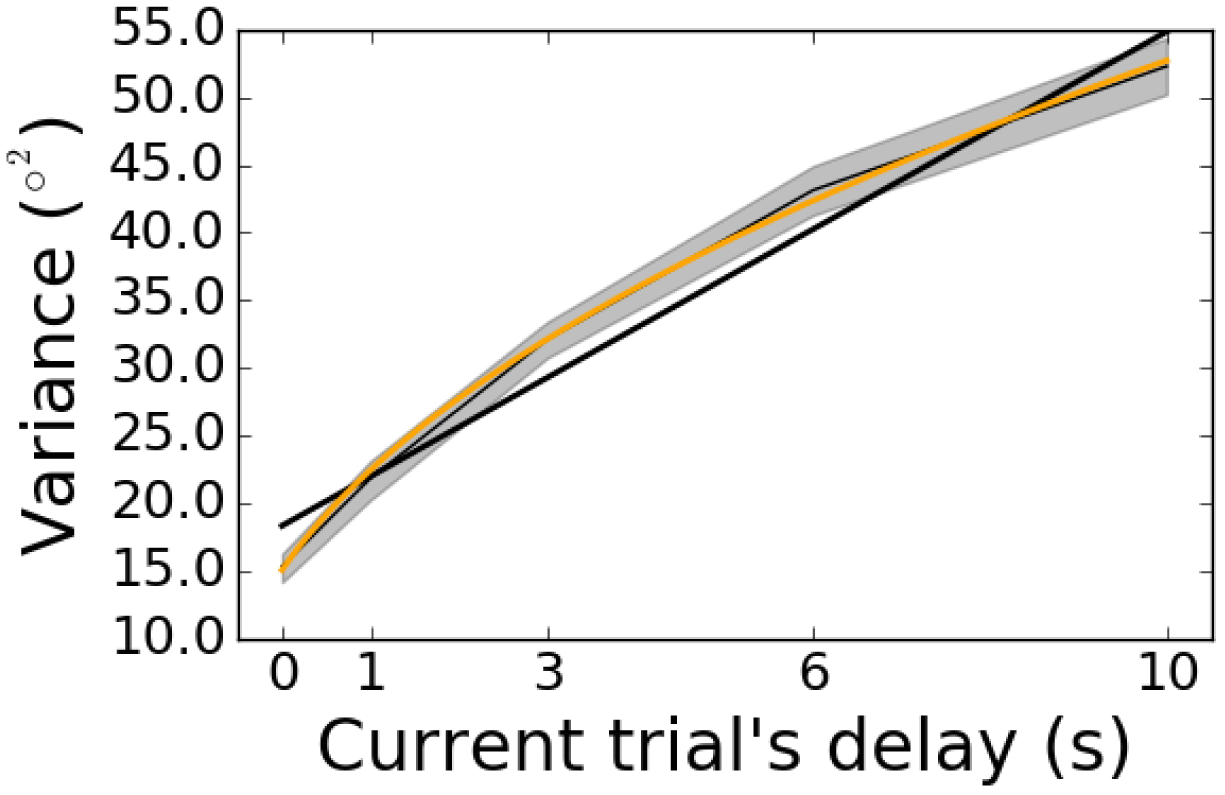
Variance of response errors as a function of the current trial’s delay in Experiment 1. The thin black line depicts the group mean, with bootstrapped 95% confidence intervals in gray shading. The thick black line is the linear best fit, which is a mismatch to the sublinear increase of variance with delay. A better fit is achieved with a power law (modified to allow for non-zero variance in the 0-s condition), depicted in orange. The power law is *y* ∼ (*x*+*t*)^β^, with *β* = 0.47.

The large sample of participants from whom we collected data enabled us to assess the nature and range of individual differences in the pattern of trial-history effects that we observed at the group level. Adaptation and serial dependence are subtle effects – just a few degrees in magnitude at their peaks – whose tuning can be measured accurately only with many trials. Hence, we were unable to reliably estimate confidence intervals or compute significance for each delay length for each individual subject. Instead, we collapsed over delay conditions for the purposes of evaluating which (if any) trial-history effect dominated across time points in perception and working memory in each participant. The results are displayed in Figure 4. Participants fell into three categories. Seven subjects showed evidence of strong repulsive adaptation that dominated across time points (all *p <* 0.05, permutation tests; all bootstrapped 95% confidence intervals < 0). One of these, whose repulsive bias was strongest (peak-to-peak = −5.08°), is presented in Figure 4B. Another 11 subjects had data that showed weak and/or noisy variation as a function of the previous stimulus’ location (all *n.s.).* The remaining majority of subjects *(n =* 20) displayed visibly apparent and statistically significant attractive serial dependence (all *p <* 0.05; all bootstrapped 95% confidence intervals > 0). However, among these, there was noticeable diversity in the tuning of the effect. Figure 4C shows the tuning over stimulus differences for one subject with a low-amplitude (peak-to-peak = 1.57°) and narrow attractive serial dependence surrounded by negative “peripheral bumps”^9^ (where the bias changes direction to repulsion when consecutive stimuli are far apart). In contrast, the participant highlighted in Figure 4D evinced the canonical shape of the effect – a broad spread of the attractive effect (peak-to-peak = 5.11°) and less prominent peripheral bumps.

**Figure 4.**
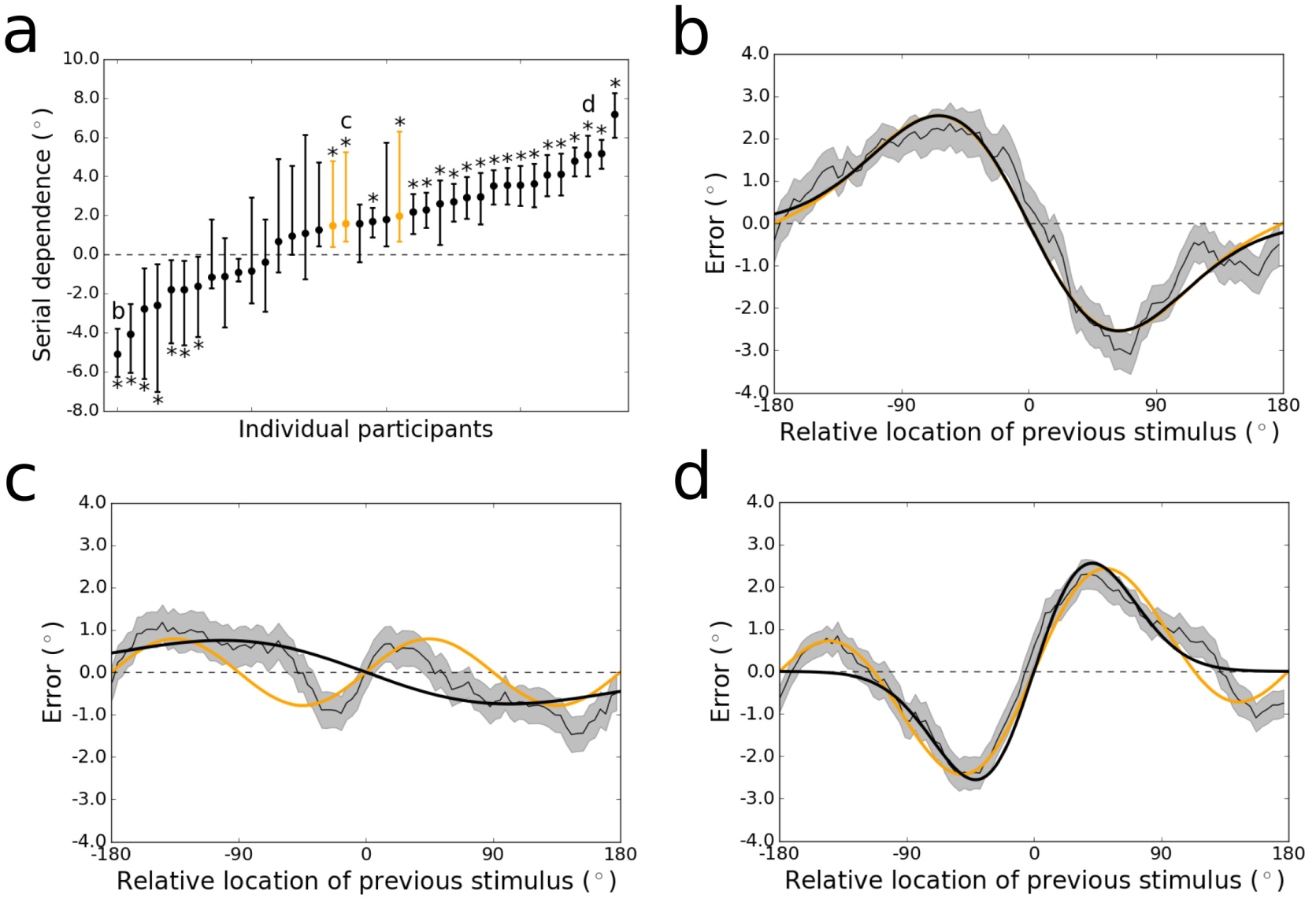
(A) Magnitudes of serial dependence observed for the individual participants tested in Experiment 1. For all but three individuals, serial dependence was measured as the peak-to-peak of the DoG fit to the data. The DoG was a qualitatively poor fit to the remaining participants (e.g., Fig. 4C), due to the prominence of peripheral bumps in their serial dependence tuning functions, which the DoG cannot capture. (The term “peripheral bumps” refers to repulsion at large differences between consecutive stimuli, in the same condition in which attraction occurs at small differences.) These participants are colored orange in this plot. Letters in this plot refer to the subfigures that follow. Significant negative adaptation and positive serial dependence (*p* < 0.05, permutation tests) are labeled with asterisks. Error bars are bootstrapped 95% confidence intervals. (B) Tuning of sensory adaptation across all possible angular differences between the current and previous stimulus, for the subject whose adaptation was strongest (peak-to-peak = −5.08^◦^. The thin black line represents the group moving average of response errors, with the standard error in gray shading, and the thick black line is the best-fitting DoG curve, which fits the data as well as the Clifford model (in orange). (C) Tuning of serial dependence for a subject with a non-canonical pattern of the effect (peak-to-peak = 1.57^◦^). The peripheral bumps are high in amplitude and wide relative to the narrow central attractive bias. The Clifford model in orange captures the positive and negative peaks of the effect well (even while the widths are misestimated), whereas the DoG mischaracterizes the bias as adaptation (negative peak-to-peak). (D) Tuning of serial dependence for a subject with strong, canonical serial dependence (peak-to-peak = 5.11^◦^). Here, the central peaks of serial dependence are wider and higher-amplitude than the peripheral bumps, and both the DoG (in black) and Clifford model (in orange) capture the magnitude of the effect well, though the DoG misses the peripheral repulsion.

**Figure 5.**
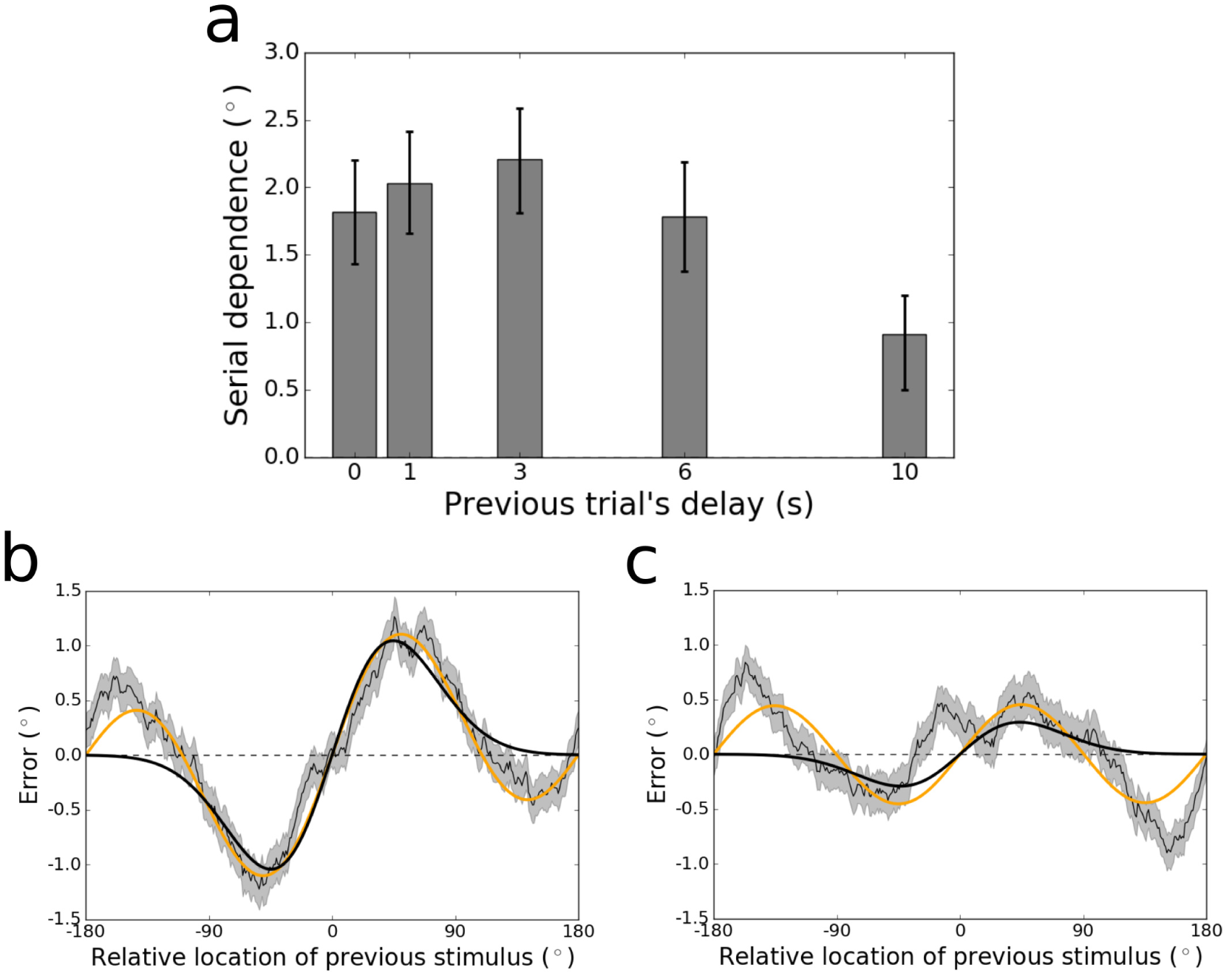
(A) Peak-to-peak of serial dependence in the group data for each length of the previous trial’s delay tested in Experiment 1. The peak-to-peak was calculated using a least squares fit of the Clifford tuning function to the data. Statistics could not be computed reliably using the DoG function due to its inability to capture the “peripheral bumps” of serial dependence, which were prominent when the data were sorted by the length of the previous delay period (see Supp. Fig. 1). Error bars represent bootstrapped 95% confidence intervals. Serial dependence is constant between 0 and 6 s, but then drops in magnitude between 6 and 10 s. (B) Tuning of serial dependence across all possible angular differences between the current and previous stimulus, for responses that followed trials with a delay length of 3 s. The thin black line represents the group moving average of response errors, with the standard error in gray shading, and the thick orange line is the best-fitting Clifford curve (2.21^◦^ peak-to-peak). The DoG fit (thick black line) misses the large-amplitude peripheral bumps at the extremes of the x-axis in this plot. (C) Tuning of serial dependence when the preceding trial’s delay period was 10 s. Here, the peak-to-peak of the Clifford fit is 0.91^◦^. The attractive peaks of serial dependence in this condition are clearly reduced relative to Figure ??B, but the peripheral bumps are just as prominent. The best DoG fit reasonably approximates the magnitude of the attractive effect, but its failure to account for the peripheral bumps causes resampling and permutation statistics to be unstable (Supp. Fig. 1).

**Figure 6.**
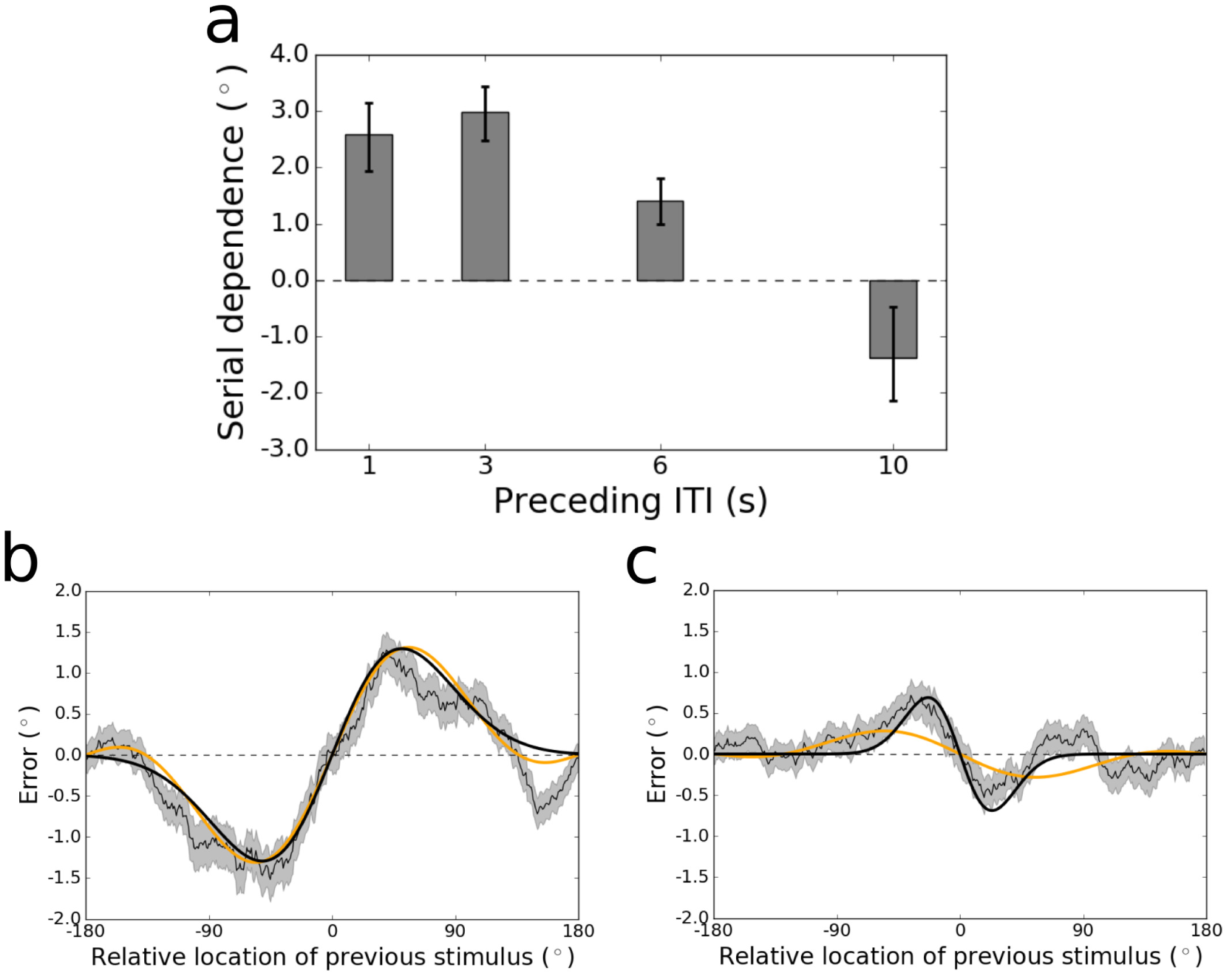
(A) Peak-to-peak of serial dependence in the group data for each ITI tested in Experiment 2. The peak-to-peak was calculated using a least squares fit of the DoG tuning function to the data. Error bars represent bootstrapped 95% confidence intervals. Serial dependence decreases in magnitude as the ITI lengthens, and then flips to significant repulsive sensory adaptation at an ITI of 10 s. (B) Tuning of serial dependence across all possible angular differences between the current and previous stimulus, for the 1-s ITI condition. The thin black line represents the group moving average of response errors, with the standard error in gray shading, and the thick black line is the best-fitting DoG curve (2.59^◦^ peak-to-peak). In orange is the best fit of the Clifford model, which captures the amplitude of the effect as well as the DoG. (C) Tuning of sensory adaptation for the 10-s ITI condition. Here, the peak-to-peak of the DoG fit is −1.38^◦^. The Clifford model cannot conform to the narrow width of the effect in this condition and underestimates its magnitude.

The time course of serial dependence we observed at the group level over the current trial’s delay period was not reproduced when trials were sorted based on the previous trial’s delay period. For each of the possible preceding delay period lengths, serial dependence in the current trial’s response was significantly greater than zero (all *p <* 0.01, group permutation tests; Fig. ??A). Between 0 and 6 s (of delay on the previous trial), serial dependence varied little (all comparisons *n.s.,* group permutation tests; minimum peak-to-peak at 6 s = 1.78°; maximum peak-to-peak at 3 s = 2.21°; all bootstrapped 95% confidence intervals overlapping). However, when the previous delay was as long as 10 s, serial dependence was significantly reduced relative to each of the other delay lengths (all *p <* 0.005; peak-to-peak at 10 s = 0.91°; bootstrapped 95% confidence interval = [0.50°, 1.20°]). Tuning for the conditions with the strongest (3 s) and weakest (10 s) serial dependence are displayed in Figures ??B and C, respectively. This set of findings is partially consistent with results from another study that tested a narrower range of delays (0.8-3.2 s) in non-human primates^16^. In this earlier study, it was found that the amplitude of serial dependence remained constant over this range (of previous delay length). Here, we extend this result to show that the previous trial’s influence does eventually decay when the previous trial’s delay is especially long.

In line with past work on the sources of error in working memory^9,33^, our findings suggest that the increasing serial dependence over longer delays (in the current trial) reflects its association with mnemonic processes rather than perceptual and motor processes that were held constant in our experiment. Research over the last decade has yielded several mathematical models designed to isolate distinct sources of error in working memory^34,35, 37–41^. None of these include parameters for the proactive interference that serial dependence represents^32^. Also, the only substantive difference between the models is their characterization of noise in the distribution of behavioral responses. As a form of systematic error, serial dependence is separable from noise, and so can be incorporated into any of these models without changing their definitions or differences. The simplest model (sometimes called the “equal precision” model^37,39^) fits random error with a single von Mises distribution^34^. In contrast, the “variable precision” model assumes the standard deviation parameter of the von Mises varies from trial to trial according to a gamma distribution^37,38^. A third model explicitly regards the precision of working memory as arising from noise in Poisson-distributed spike trains of individual neurons^40,41^. Errors in this model are distributed according to a von Mises random walk^41^. We will refer to these three working memory models as EP (equal precision), VP (variable precision), and VMRW (von Mises random walk).

As a first pass, we fit each of these models to the behavioral data from Experiment 1. Model comparison on the basis of the corrected Akaike Information Criterion (AICc) revealed that the VMRW model fit the data better than the VP model (ΔAICc = 21.7 ±5.2), which fit the data better than the EP model (ΔAICc = 18.4 ±5.5). This ordering is consistent with published comparisons of the VP and EP models using behavioral data from other working memory tasks^37–39^. However, a previous study that compared the VMRW model with the VP model found no consistent difference between the goodness of their fits using this metric^40^. That comparison was based on data from a total of eight participants, relative to 38 in our analysis (with similar numbers of trials completed by each participant in both studies). Hence, relative to our study, the earlier analysis may have been underpowered to detect differences between these two models, which predict similar deviations from normality in working memory error distributions.

Next, we created a hybrid model that incorporates serial dependence into the mean of the VMRW distribution – sliding the mean clockwise or counterclockwise on each trial by the magnitude dictated by the tuning of the history effect (see Methods). This hybrid model significantly outperforms the base VMRW model (ΔAICc = 29.2±7.9). However, this result on its own falls short of confirming that the serial dependence tuning function is needed to quantify the influence of the history effect on each trial. To verify that inclusion of the tuning curve visible in Figures 2 and 4-?? is needed for the improvement in fit, we developed an alternative hybrid “memory confusion”^2^ model that takes trial history into account in a different way. In this model, it is assumed that on a subset of trials, subjects simply mix up which stimulus belongs to the current trial and report the previous trial’s location when probed (analogous to a “swap”^36,43^ over time rather than space). This “memory confusion” model provides no benefit above the base VMRW model and makes it worse, due to the addition of parameters that capture little variance (ΔAICc = —7.7 ± 2.7). The addition of the serial dependence tuning function to all three of the base models does not change the order of performance among them: the improved VMRW model outperforms the improved VP model (ΔAICc = 12.5 ± 4.2), which outperforms the improved EP model (ΔAICc = 14.3 ± 5.5).

### Experiment 2: Manipulation of baseline interval between trials

It is possible that the delay manipulation in Experiment 1 confounded two variables: (1) the time for which subjects must hold the current item in memory and (2) the time that has elapsed since the behavioral response on the previous trial, before the current trial’s response. To demonstrate that the time course of serial dependence we observed (Fig. 2A) corresponds to mnemonic processes and not the simple passage of time, we conducted a second experiment in which the inter-trial interval (ITI) varied randomly among 1, 3, 6, and 10 s. The delay in this new task was held constant at 3 s. In all other respects, the tasks for the two experiments were identical.

Collapsing across ITIs, we identified serial dependence in the group dataset significantly greater than zero *(p <* 10^−4^, group permutation test; peak-to-peak = 1.62°; bootstrapped 95% confidence interval = [1.38°, 1.86°]). As for Experiment 1, there was no bias in the data in the direction of the stimulus on the upcoming trial *(n.s.,* group permutation test; peak-to-peak = 0.23°; bootstrapped 95% confidence interval = [−0.15°, 0.80°]), an important control^10,14^. This pair of results replicates our finding from Experiment 1 of serial dependence in this spatial delayed response task, using an independent dataset.

Next we examined each of the ITIs individually. The magnitude of serial dependence across ITIs from 1-10 s is plotted in Figure ??A. The magnitude of serial dependence decreases gradually during the interval between trials, marginally from 3-6 s *(p =* 0.01, group permutation test, Bonferroni-corrected *α =* 0.008; lower bound of bootstrapped 95% confidence interval at 3 s = 2.48°; upper bound of confidence interval at 6 s = 1.81°) and significantly from 6-10 s *(p <* 10^−4^; lower bound of bootstrapped 95% confidence interval at 6 s = 1.00°; upper bound of confidence interval at 10 s = 0.48°). The difference in serial dependence between the 1-s (Fig. ??B) and 3-s ITIs was statistically non-significant. The slope of this time course is opposite that obtained in Experiment 1, strengthening our conclusion that increased serial dependence with increased delay length is due to the prolongation of memory demands rather than the mere passage of time. For the largest ITI (10 s), participants’ responses on the trial after the ITI were repelled away from the preceding trial’s stimulus, an effect consistent with sensory adaptation *(p =* 0.006; peak-to-peak = -1.38°; bootstrapped 95% confidence interval = [−2.15°, −0.48°]; Fig. ??C). In contrast, for every other ITI tested, serial dependence was significantly greater than zero (all *p ≤* 10^−3^, group permutation tests).

## Discussion

In everyday visual experience, humans rely not just on moment-to-moment perception but also on continued maintenance of information in working memory to navigate their environments and accomplish tasks. While there is much evidence to suggest that working memory recruits the same cortical areas active during sensory perception^49–57^, remembered visual content differs in quality^33,47,58^ – and potentially representational format^59,60^ – from feedforward signals driven by the presence of an external stimulus. Both behavioral data^33,47,58^ and computational theory^48^ have implied that passage of visual percepts into memory makes them less precise. This past work has also claimed that mnemonic processes do not attach to percepts any accumulating systematic bias – just random noise due to drift and/or decay^33,47,48,58^. With the experiments reported here, we provide new evidence to disconfirm this view. Serial dependence – a systematic bias in the direction of the preceding trial’s stimulus – is absent from percepts until the working memory system is engaged. Our demonstration of repulsive adaptation – with no attractive serial dependence – in the perception condition extends previous work^9^ by showing that this oppositely valenced effect that precedes working memory does not require that subjects make a comparison between two simultaneously presented stimuli^9^ – adaptation occurs in the context of the same delayed response task that yields serial dependence when memory demands are increased. To our knowledge, this is the first report of sensory adaptation in a spatial delayed response task, though adaptation for other visual features is well established^20–24^.

By testing a wider range of delays between stimulus and response than used in previous studies^9,16,17^, we were able to chart the time course of serial dependence in visual working memory. This technique – of probing participants to report the contents of memory at variable points in time after stimulus offset – is common in visual psychophysics^61^. It has revealed how information passing through the visual system progresses from a rich perceptual code to a more impoverished mnemonic one. For a few hundred milliseconds after visual input ceases, a great deal of perceptual detail is still accessible to the observer in iconic memory – a form of storage intermediate between perception and working memory^62^. After that, within one second of delay, capacity-limited, distraction-resistant working memory comes online in parallel with a larger-capacity system that is vulnerable to distraction – fragile memory^44,45,63–66^. Our experiments demonstrate that the residual sensory trace associated with iconic memory is free of serial dependence – though it does carry the opposite, repulsive bias associated with sensory adaptation. The attractive bias arises slowly in the later short-term memory systems, but asymptotes before long-term storage processes are engaged (at approximately 20 seconds of delay^47^). Future research may resolve with finer resolution the exact moment at which serial dependence appears and whether it is most strongly associated with fragile or distraction-resistant stages of working memory. (Consistent with most work in this area^67,68^, we have tended to use the term “working memory” as a shorthand for both of these systems.)

Beyond demonstrating that serial dependence accumulates for longer in working memory than previous studies have indicated^9,16,17^, we have taken strides to integrate this phenomenon into the study of working memory in ways it has not been before^32^. Specifically, we have made concrete, formal improvements to prominent mathematical models designed to characterize the psychological architecture of working memory. The provision of terms for serial dependence to these models both (1) allows them to capture more variance in behavioral data and (2) ensures that the variance associated with the temporal smoothing operation of serial dependence does not distort estimates of the models’ other parameters. Claims that have been made about the nature of decay rates in working memory without consideration of trial-history biases must now be ree¨valuated. For example, one study that modeled behavioral responses following different delay period lengths concluded that maintained representations are susceptible to spontaneous complete erasure from working memory as the delay length increases (measured as an increase in guess rates), but not to subtle degradations in precision (measured with the *κ* parameter of a variation on the EP model)^42^. However, because this study ignored potential serial dependencies in the data, as well as alternative models of noise (e.g., VP and VMRW), the validity of this conclusion is unclear. It is impossible to address claims about total loss of information from memory with our data, because guess rates in our simple one-item spatial task were near zero. In the future, however, the hybrid models we have developed that incorporate serial dependence may help elucidate the nature of working memory storage in more difficult multi-item tasks. To what extent serial dependencies occur when multiple items are held in mind at once is an open question that the hybrid models we have validated can help answer.

Our experiments have filled other gaps in the field’s understanding of the temporal properties of serial dependence. We have determined the approximate duration for which this effect persists between trials. At least in spatial working memory, the attractive bias disappears within ten seconds after the end of each trial, and is replaced by (or exposes a persistent) low-amplitude adaptation. This constrains possible neural theories of serial dependence – viable mechanisms must have time constants on the order of 10 s, which rules out especially short-term (e.g., synaptic facilitation) and long-term (e.g., long-term potentiation) forms of plasticity. Previous attempts to measure the washout period of serial dependence in humans have used a short, fixed ITI, preventing the measurement of pure time in the absence of intervening trials^1,14^. One experiment using non-human primate subjects did report a decrease in serial dependence between 2 and 7 s of ITI^16^. Over this range, the effect remained above baseline for two of three subjects, and no crossover to adaptation was observed. We have also shown that serial dependence weakens when the stimulus on the previous trial is maintained for as long as 10 s. It is possible that the neural code changes abruptly around this time point: for example, elevated neural firing keeping the representation active may begin to fail spontaneously (as happens in some neural-network models^?^), leaving the representation in an “activity silent” state supported by short-term plasticity^?, ?^. Exponential decay of this synaptic trace (without continued active firing to keep it in place) may explain the reduced influence on responses on the subsequent trial.

Reframing serial dependence as a phenomenon of working memory rather than perception does not change the theories that have been put forth about its functional importance^32^. Thus, it remains an important mechanism for stabilizing representations against interruptions in visibility^1,8^. The contents of working memory track the focus of attention^25–31^, which, during the execution of a single goal, can remain the same for several seconds, even as the raw visual input that impinges on the retina fluctuates due to saccades, occlusion, and changes in lighting. Hence, temporal autocorrelation in visual working memory is potentially even higher than it is in visual scenes (and perception). If true, this would explain why serial dependence may have evolved in working memory as opposed to perceptual circuits – more autocorrelation enhances the ability of temporal smoothing to limit the influence of noise and boost signal. Moreover, the offloading of attractive serial dependence to memory systems may accord perceptual systems enhanced capacity to specialize in novelty detection, in part via adaptation. More research is needed to elucidate the ways in which serial dependence and adaptation interact, and to reveal the ecologically valid situations in which one or the other (perhaps both at the same time) enhance visual performance^32^. Such continued study should aim to clarify the mechanisms and functional consequences of the striking diversity we observed in the strength and tuning shape of these effects across individuals.

## Methods

### Participants

Fifty-five adults (34 female) from the UC Berkeley community were recruited to participate in this study. Thirty-five of these individuals completed Experiment 1 only, fourteen completed Experiment 2 only and six completed both experiments. All aspects of data collection and analysis were conducted in accordance with guidelines approved by the Committee for the Protection of Human Subjects at UC Berkeley. Informed consent was obtained from all subjects, and they were compensated monetarily for their time.

### Experimental Procedures

Participants completed the protocol in a soundproof, dimly lit testing room. For both experiments, they completed a spatial delayed response task, depicted in Figure 1 (adapted from^46^). The task was programmed in MATLAB using the Psychophysics Toolbox^69^ (version 3) and run on a Mac mini (OSX El Capitan 10.11). For eight subjects in Experiment 1, a 17-in monitor was used with a screen resolution of 1280 X 1024 pixels. The remaining sessions were run with a 23-in monitor, 1920 X 1080 pixels. All participants were seated such that their eyes were approximately 60 cm from the center of the testing display.

The stages of the generic task used for both experiments are as follows (with angle measurements reported in degrees of visual angle). Each trial began with the presentation of a black circle for 1 s at a random polar angle from fixation, with eccentricity fixed at 12°. The circle’s diameter was 1°. All stimuli were displayed against a gray background. Participants were instructed to fixate a central black square – which spanned 0.5° X 0.5° – whenever it was on the screen (all stages of the task aside from the response period). In Experiment 1, participants remembered the location of the presented circle for a delay that varied randomly from trial to trial (0, 1,3,6, or 10 s). The delay was always 3 s in Experiment 2. At the end of the delay, the fixation square was replaced with the mouse cursor (at the exact center of the screen), and participants indicated the location in mind by moving the cursor to that location and clicking once. No feedback was given. Errors were measured in degrees of polar angle. In Experiment 1, a 1-s ITI followed the response period, before the start of the next trial. The ITI varied randomly from trial to trial in Experiment 2 (1, 3, 6, or 10 s). Each participant completed 1,000 trials (200 per delay) in Experiment 1, divided into 40 blocks over the course of one or two experimental sessions. All but two participants completed 1,008 trials (252 per ITI) in Experiment 2. The remaining two participants completed 999 and 1,017 trials, respectively.

### Data Analysis

The data were analyzed using Python, MATLAB, and shell scripts. All code written for this study is available upon request.

Before model fitting for trial-history effects, the data were submitted to preprocessing. First, trials with responses that were within 5° of visual angle of the origin were dropped, as were trials with responses further than three standard deviations from the participant’s mean error (<0.7% of all trials, across subjects). Next, we computed systematic directional error as the mean response for each stimulus location. This mean was then subtracted from the response on each individual trial (ignoring the location of the previous trial) to obtain the residual error that was used to characterize serial dependence. Replicating the procedure in^16^, we computed the systematic error by spatially low-pass filtering the responses as a function of stimulus location using the MATLAB function `loess`. Finally, to ensure that our analyses were restricted to data from participants who performed the task correctly, we removed those with noticeably poor performance. Specifically, we removed subjects with an overall mean absolute error greater than 10° of polar angle. This criterion, though arbitrary, removed only subjects with qualitatively noisy error histograms while retaining those whose errors were roughly normally distributed around the correct value (the expected pattern). Only three subjects in Experiment 1 failed to pass this criterion (mean absolute error 37.3 ± 19.5 for these three compared to 4.7 ± 0.2 for the others). Data from two subjects in Experiment 2 were excluded (53.9 ± 1.8 for these compared to 4.7 ± 0.4 for the others).

Studies that have modeled the tuning of serial dependence to featural differences between past and current visual stimuli have used the derivative of Gaussian (DoG)^1,2,8,9^ (or the very similar Gabor function^16, 17^). There is another function in the perception literature, developed by Clifford and colleagues^20^, that has been used to model sensory adaptation – and that therefore fits serial dependence readily (when multiplied by -1). Overall, these functions fit the data from our Experiment 1 equivalently well (collapsing over delays, ΔAICc = 0.8 ± 1.0, favoring DoG over Clifford), and this equivalence also holds for Experiment 2 (collapsing over ITIs, ΔAICc = 1.1 ± 1.0, favoring DoG over Clifford). However, we noted significant differences between the Clifford and DoG models in certain conditions. Occasionally, the attractive bias of serial dependence is accompanied by a repulsion effect (“peripheral bumps”^9^) when previous visual input is close to maximally different from the input on the current trial (the extremes of the x-axis in Fig. 2C). The DoG cannot account for this reversal of the response bias, so when it is prominent in the data, the best fit of the DoG tends to mischaracterize the true effect size (e.g., 4C). The Clifford model is a combination of sinusoids of different frequencies designed to capture the peripheral bumps when they appear^20^. However, when the trial-history effect is narrow over stimulus differences and there are no peripheral bumps, the Clifford model tends to fail (e.g., Fig. 2A). This is because the Clifford model – unlike the DoG – does not have an independent width parameter; shrinking the central width of the Clifford fit requires that the peripheral bumps be increased. To be consistent with previous literature on serial dependence, we use the DoG for all analyses, except in cases where it provides a poor fit to the data, in which case we use the Clifford model (as noted below). The mathematical definitions of both models are reported next.

In this study, differences between past and current visual input ranged between -180 and 180° of polar angle (a complete circle). The DoG is defined as

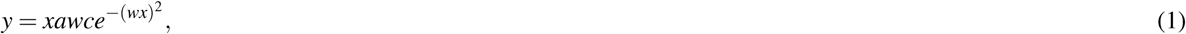

where *y* is the signed error, *x* is the relative angle of the previous trial, *a* is the amplitude of the curve peaks, *w* is the width of the curve, and *c* is the constant 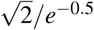.

The Clifford model is stated as follows:

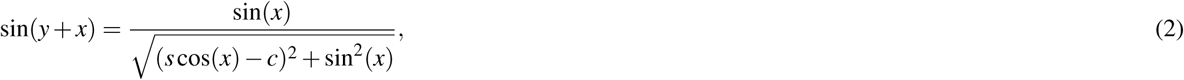

where *s* is a scaling parameter and *c* is a centering parameter.

We used the scipy^70^ function `least_squares` (in the `optimize` module) to find the values of a and *w*, in the case of the DoG, or *c* and *s,* in the case of the Clifford model, that minimized the difference, for each x, between the estimated *y* and the subject’s actual error. Across all values of x, we take the magnitude of serial dependence (or adaptation) to be the peak-to-peak of *y*(*x*), with the sign adjusted to match the direction of the effect (see Fig. 2C).

To determine whether the magnitude of serial dependence was significantly greater than zero, or greater in one condition than in another, we submitted the data to permutation testing at the group level^2,9^. Specifically, we shuffled the values of x (current trial’s location relative to the previous trial’s) while leaving in place the corresponding errors. We then fit the DoG to the shuffled dataset. This process was repeated 10,000 times. The *p*-values we report are the proportion of permutations that led to equal or higher values for the peak-to-peak of the function fit than the one estimated for the unshuffled data. In the case of a comparison between conditions, we subtracted the null peak-to-peaks for one condition from those for the other, and report the proportion of these differences that had equal or higher values than the empirical difference. The criterion for significance was Bonferroni-corrected for each family of tests.

We computed bootstrapped confidence intervals as follows^2,9^: We resampled the data with replacement 10,000 times. We then fit the DoG to each resampled dataset. This yielded a distribution of peak-to-peak values from which we selected the boundaries of the 95% confidence interval – separately for each delay and ITI condition.

In our statistical analyses of group data, only one condition – the 10-s condition in the analysis of the previous trial’s delay length for Experiment 1 – could not be fit reliably using the DoG, due to large peripheral bumps (Fig. ??C; Supp. Fig. 1). Hence, for this analysis we used the Clifford model, which estimated the peak-to-peak reliably. In our plot of individual subjects’ data (Fig. 4A), the DoG fit (with bootstrapped confidence intervals) is reported for all but three subjects, for whom the Clifford model was a qualitatively superior fit. These three subjects are labeled in the figure, and one is highlighted in Figure 4C.

Three base mathematical models of working memory – EP^34^, VP^37,38^, and VMRW^40,41^ – were fit to our behavioral data, as described in the Results. Model fitting was done using the MATLAB function `fminsearch`, separately for each delay condition. EP is defined as

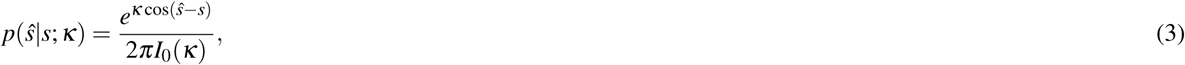

the von Mises probability density function. Here, *§* is each trial’s response, *s* the corresponding stimulus, and *I*_0_ the modified Bessel function of the first kind, order zero. The concentration parameter, *κ,* is a measure of response precision, spanning all trials, and is the model’s one free parameter for fitting.

In VP, precision is drawn anew for each trial from a gamma (*γ*) distribution with mean *J* and scale parameter *τ* (the model’s free parameters). Built from EP, this gives

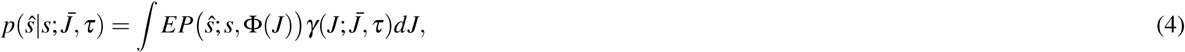

where EP’s concentration parameter *κ* is a function of *J* – here expressed as Φ(*J*) – and *J* is formally defined as Fisher information. The relation *κ* = Φ(*J*) can be computed numerically^37^, but this relation was not needed for the present analyses. The integral in Equation 4 has no analytical expression and so is approximated using Monte Carlo simulations^37,39^.

Finally, in VMRW, noise in working memory is distributed according to a von Mises random walk, as derived from a population coding model of cortex^41^. Specifically, behavioral errors for a random walk of length *r* are von Mises distributed:

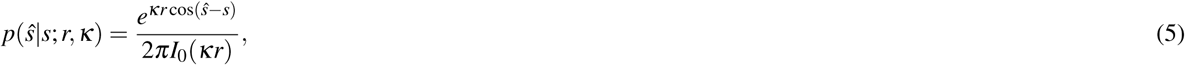

where the distribution of r for *m* walk steps is

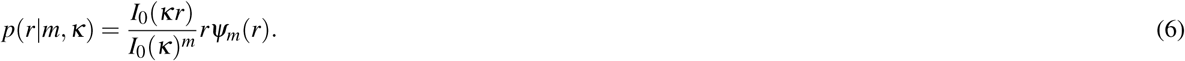

Here, *rΨ_m_*(*r*) is the probability density function for a uniform random walk of length *r* and number of steps *m*. The variable *m* is itself Poisson-distributed, with expected value *ξ* For additional equations and a full derivation, including the neural interpretation of these variables, see^41^. In order to fit this model to data, we approximated the density *Ψ_m_*(*r*) via Monte Carlo simulation. The free variables for fitting are *κ* (the concentration parameter) and *ξ,* which corresponds to gain.

We added terms to these base models to capture temporal smoothing in the data in the form of serial dependence (or adaptation). In particular, we allowed the mean of each model’s probability density function to vary on a trial-by-trial basis, as a function of the location of the previous trial’s stimulus. Given a particular difference in location between the current and previous trial’s stimuli, the mean shift was set to be the value of the DoG model fit to the data at that point. (That is, in visual terms, the input to the model was a point on the x-axis in Figure 2C, for example, and the output mean shift was the DoG function’s value on the y-axis.) This procedure added two additional variables to each of the base models – *a* and *w.*

As an alternative to the base models with the serial dependence expansion, we made alternative models that account for trial history by assuming that participants, on a subset of trials, confuse which stimulus was presented most recently and report the wrong item when probed. This alternative similarly allowed the mean of the base probability density functions to shift, depending on the difference between the previous trial’s location and the current one, without altering their shape or width. This “swap over time” model is defined as^36^

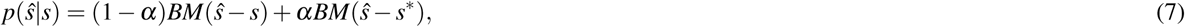

where *BM* is a base model, *α* (an additional free parameter) sets the frequency of swaps, and *s\** is the stimulus location for the previous trial.

Within each model, we used a separate set of parameters for each memory delay length, and formally compared the fits of different models using the Akaike Information Criterion (as recommended in^39^), with the standard correction for finite sample sizes (AICc). AICc values were averaged across subjects for these comparisons.

## Author Contributions

D.P.B. and M.D. designed the experiment and wrote the paper. D.P.B. prepared the stimuli. D.P.B. and J.J.S. tested the subjects and analysed the results.

## References

1. Fischer, J. & Whitney, D. Serial dependence in visual perception. Nature Neuroscience 17, 738–743 (2014).

2. Liberman, A., Fischer, J. & Whitney, D. Serial dependence in the perception of faces. Current Biology 24, 2569–2574 (2014).

3. Xia, Y., Leib, A. Y. & Whitney, D. Serial dependence in the perception of attractiveness. Journal of vision 16, 28–28 (2016).

4. Huang, J. & Sekuler, R. Distortions in recall from visual memory: Two classes of attractors at work. Journal of Vision 10, 24–24 (2010).

5. Kondo, A., Takahashi, K. & Watanabe, K. Sequential effects in face-attractiveness judgment. Perception 41, 43–49 (2012).

6. Ward, L. M. & Lockhead, G. R. Response system processes in absolute judgment. Attention, Perception, & Psychophysics 9, 73–78 (1971).

7. Petzold, P. & Haubensak, G Higher order sequential effects in psychophysical judgments. Attention, Perception, & Psychophysics 63, 969–978 (2001).

8. Liberman, A., Zhang, K. & Whitney, D. Serial dependence promotes object stability during occlusion. Journal of Vision 16, 16–16 (2016).

9. Fritsche, M., Mostert, P. & de Lange, F P. Opposite effects of recent history on perception and decision. Current Biology (2017).

10. Cicchini, G M., Anobile, G & Burr, D. C. Compressive mapping of number to space reflects dynamic encoding mechanisms, not static logarithmic transform. Proceedings of the National Academy of Sciences 111, 7867–7872 (2014).

11. Kondo, A., Takahashi, K. & Watanabe, K. Influence of gender membership on sequential decisions of face attractiveness. Attention, Perception, & Psychophysics 75, 1347–1352 (2013).

12. Ward, L. M. Mixed-modality psychophysical scaling: Inter-and intramodality sequential dependencies as a function of lag. Attention, Perception, & Psychophysics 38, 512–522 (1985).

13. Petzold, P. & Haubensak, G The influence of category membership of stimuli on sequential effects in magnitude judgment. Perception & Psychophysics 66, 665–678 (2004).

14. Taubert, J., Alais, D. & Burr, D. Different coding strategies for the perception of stable and changeable facial attributes. Scientific Reports 6 (2016).

15. Corbett, J. E., Fischer, J. & Whitney, D. Facilitating stable representations: Serial dependence in vision. PloS One 6, e16701 (2011).

16. Papadimitriou, C, Ferdoash, A. & Snyder, L. H. Ghosts in the machine: memory interference from the previous trial. Journal of neurophysiology 113, 567–577 (2015).

17. Papadimitriou, C, White, R. L. & Snyder, L. H. Ghosts in the machine ii: Neural correlates of memory interference from the previous trial. Cerebral Cortex bhw106 (2016).

18. Dong, D. W. & Atick, J. J. Statistics of natural time-varying images. Network: Computation in Neural Systems 6, 345–358 (1995).

19. Gibson, J. J. & Radner, M. Adaptation, after-effect and contrast in the perception of tilted lines. i. quantitative studies. Journal of Experimental Psychology 20, 453 (1937).

20. Clifford, C. W., Wenderoth, P. & Spehar, B. A functional angle on some after-effects in cortical vision. Proceedings of the Royal Society of London B: Biological Sciences 267, 1705–1710 (2000).

21. Webster, M. A. Visual adaptation. Annual review of vision science 1, 547–567 (2015).

22. Webster, M. A. & MacLeod, D. I. Visual adaptation and face perception. Philosophical Transactions of the Royal Society B: Biological Sciences 366, 1702–1725 (2011).

23. Webster, M. A. Adaptation and visual coding. Journal of vision 11, 3–3 (2011).

24. Clifford, C. W. et al. Visual adaptation: Neural, psychological and computational aspects. Vision research 47, 3125–3131 (2007).

25. Kiyonaga, A. & Egner, T. Working memory as internal attention: toward an integrative account of internal and external selection processes. Psychonomic bulletin & review 20, 228–242 (2013).

26. Chun, M. M. Visual working memory as visual attention sustained internally over time. Neuropsychologia 49, 1407–1409 (2011).

27. Chun, M. M. & Johnson, M. K. Memory: enduring traces of perceptual and reflective attention. Neuron 72, 520–535 (2011).

28. Gazzaley, A. & Nobre, A. C. Top-down modulation: bridging selective attention and working memory. Trends in cognitive sciences 16, 129–135 (2012).

29. Awh, E. & Jonides, J. Overlapping mechanisms of attention and spatial working memory. Trends in cognitive sciences 5, 119–126 (2001).

30. Awh, E., Vogel, E. & Oh, S.-H. Interactions between attention and working memory. Neuroscience 139, 201–208 (2006).

31. Myers, N. E., Stokes, M. G. & Nobre, A. C. Prioritizing information during working memory: Beyond sustained internal attention. Trends in Cognitive Sciences (2017).

32. Kiyonaga, A., Scimeca, J. M., Bliss, D. P. & Whitney, D. Serial dependence across perception, attention, and memory. Trends in Cognitive Sciences (2017).

33. White, J. M., Sparks, D. L. & Stanford, T. R. Saccades to remembered target locations: an analysis of systematic and variable errors. Vision research 34, 79–92 (1994).

34. Wilken, P. & Ma, W. J. A detection theory account of change detection. Journal of vision 4, 11–11 (2004).

35. Zhang, W. & Luck, S. J. Discrete fixed-resolution representations in visual working memory. Nature 453, 233–235 (2008).

36. Bays, P. M., Catalao, R. F. & Husain, M. The precision of visual working memory is set by allocation of a shared resource. Journal of vision 9, 7–7 (2009).

37. van den Berg, R., Shin, H., Chou, W.-C., George, R. & Ma, W. J. Variability in encoding precision accounts for visual short-term memory limitations. Proceedings of the National Academy of Sciences 109, 8780–8785 (2012).

38. Fougnie, D., Suchow, J. W. & Alvarez, G. A. Variability in the quality of visual working memory. Nature communications 3, 1229 (2012).

39. van den Berg, R., Awh, E. & Ma, W. J. Factorial comparison of working memory models. Psychological Review 121, 124 (2014).

40. Bays, P. M. Noise in neural populations accounts for errors in working memory. Journal of Neuroscience 34, 3632–3645 (2014).

41. Bays, P. M. A signature of neural coding at human perceptual limits. Journal of Vision 16, 4–4 (2016).

42. Zhang, W. & Luck, S. J. Sudden death and gradual decay in visual working memory. Psychological science 20, 423–428 (2009).

43. Bays, P. M. Evaluating and excluding swap errors in analogue tests of working memory. Scientific reports 6, 19203 (2016).

44. Sligte, I. G., Scholte, H. S. & Lamme, V. A. Are there multiple visual short-term memory stores? PLOS one 3, e1699 (2008).

45. Sligte, I. G., Vandenbroucke, A. R., Scholte, H. S. & Lamme, V. A. Detailed sensory memory, sloppy working memory. Frontiers in psychology 1, 175 (2010).

46. Almeida, R., Barbosa, J. & Compte, A. Neural circuit basis of visuo-spatial working memory precision: a computational and behavioral study. Journal of neurophysiology 114, 1806–1818 (2015).

47. Ploner, C. J., Gaymard, B., Rivaud, S., Agid, Y. & Pierrot-Deseilligny, C. Temporal limits of spatial working memory in humans. European Journal of Neuroscience 10, 794–797 (1998).

48. Compte, A., Brunel, N., Goldman-Rakic, P. S. & Wang, X.-J. Synaptic mechanisms and network dynamics underlying spatial working memory in a cortical network model. Cerebral Cortex 10, 910–923 (2000).

49. Harrison, S. A. & Tong, F. Decoding reveals the contents of visual working memory in early visual areas. Nature 458, 632–635 (2009).

50. Serences, J. T., Ester, E. F., Vogel, E. K. & Awh, E. Stimulus-specific delay activity in human primary visual cortex. Psychological science 20, 207–214 (2009).

51. Postle, B. R. Working memory as an emergent property of the mind and brain. Neuroscience 139, 23–38 (2006).

52. D’Esposito, M. From cognitive to neural models of working memory. Philosophical Transactions of the Royal Society B: Biological Sciences 362, 761–772 (2007).

53. D’Esposito, M. & Postle, B. R. The cognitive neuroscience of working memory. Annual review of psychology 66, 115–142 (2015).

54. Pasternak, T. & Greenlee, M. W. Working memory in primate sensory systems. Nature Reviews Neuroscience 6, 97–107 (2005).

55. Sreenivasan, K. K., Curtis, C. E. & D’Esposito, M. Revisiting the role of persistent neural activity during working memory. Trends in cognitive sciences 18, 82–89 (2014).

56. Gabrieli, J. D. Cognitive neuroscience of human memory. Annual review of psychology 49, 87–115 (1998).

57. Fuster, J. M. Network memory. Trends in neurosciences 20, 451–459 (1997).

58. Cappiello, M. & Zhang, W. A dual-trace model for visual sensory memory. Journal of Experimental Psychology: Human Perception and Performance 42, 1903–1922 (2016).

59. Sreenivasan, K. K., Vytlacil, J. & D’Esposito, M. Distributed and dynamic storage of working memory stimulus information in extrastriate cortex. Journal of cognitive neuroscience 26, 1141–1153 (2014).

60. Meyers, E. M., Freedman, D. J., Kreiman, G., Miller, E. K. & Poggio, T. Dynamic population coding of category information in inferior temporal and prefrontal cortex. Journal of neurophysiology 100, 1407–1419 (2008).

61. Magnussen, S. Low-level memory processes in vision. Trends in neurosciences 23, 247–251 (2000).

62. Sperling, G. The information available in brief visual presentations. Psychological monographs: General and applied 74, 1–29 (1960).

63. Sligte, I. G., Wokke, M. E., Tesselaar, J. P., Scholte, H. S. & Lamme, V. A. Magnetic stimulation of the dorsolateral prefrontal cortex dissociates fragile visual short-term memory from visual working memory. Neuropsychologia 49, 1578–1588 (2011).

64. Vandenbroucke, A. R., Sligte, I. G. & Lamme, V. A. Manipulations of attention dissociate fragile visual short-term memory from visual working memory. Neuropsychologia 49, 1559–1568 (2011).

65. Pinto, Y., Sligte, I. G., Shapiro, K. L. & Lamme, V. A. Fragile visual short-term memory is an object-based and location-specific store. Psychonomic bulletin & review 20, 732–739 (2013).

66. Vandenbroucke, A. R., Sligte, I. G., de Vries, J. G., Cohen, M. X. & Lamme, V. A. Neural correlates of visual short-term memory dissociate between fragile and working memory representations. Journal of cognitive neuroscience 27, 2477–2490 (2015).

67. Ma, W. J., Husain, M. & Bays, P. M. Changing concepts of working memory. Nature neuroscience 17, 347–356 (2014).

68. Luck, S. J. & Vogel, E. K. Visual working memory capacity: from psychophysics and neurobiology to individual differences. Trends in cognitive sciences 17, 391–400 (2013).

69. Brainard, D. H. The psychophysics toolbox. Spatial vision 10, 433–436 (1997).

70. Jones, E., Oliphant, T. & Peterson, P. Scipy: Open source scientific tools for python. http://www.scipy.org/ (2001).

